# Elemental analysis of *Moringa oleifera* seeds by Laser Induced Breakdown Spectroscopy (LIBS) and its anti-cancer and anti-microbial studies

**DOI:** 10.1101/2020.04.15.042663

**Authors:** R. K. Aldakheel, S. Rehman, Firdos A. Khan, A. Mostafa, M. A. Almessiere, M. A. Gondal, A. Baykal

## Abstract

Moringa oleifera plant contains a numerous antioxidants, antibiotics and nutrients (vitamins and minerals) which makes it prospective for diverse biomedical applications. This report investigated the anti-cancerous and anti-microbial potential of the Moringa oleifera seeds (MOSs) due to the bioactive components a detail elemental analysis. MOSs in the form of pellets were used in elemental analysis. Alcoholic extraction of were utilized in anti-cancerous and anti-microbial activity.To clarify the anti-cancerous and anti-microbial potential of the (MOSs) because of the bioactive components. An elemental analysis was conducted using Laser Induced Breakdown Spectroscopy (LIBS). The GC-MS used for the validation of the LIBS outcome. Additionally, the anti-cancerous and anti-microbial activity of the MOSs was evaluated. Herein, the human colorectal carcinoma cells (HCT-116) were treated with the seeds aqueous extracts for 48 h and the cell viability plus the DNA nuclear morphology were measured via the MTT assay and DAPI staining. The cell viability of the normal human embryonic stem cells (HEK-293) was also examined after the treatment. Percentage of cell viability and inhibitory concentration (IC_50_) of both normal and treated cells were determined.The recorded LIBS signal of the MOSs revealed the existence of elements like Ca, K, Mg, P, S, Fe, Mn, Zn, Na and Se (vital for human health). The results of the MTT assay revealed a profound inhibitory action of the MOSs extracts against the HCT-116 cells growth. On top, such extracts did not affect the HEK-293 cells growth, indicating the specificity of the proposed extracts towards cancer cells escalation hindrance. The anti-microbial activity of the ethanolic MOSs extracts was tested on the S. aureus and E. coli bacteria using the Agar well diffusion assay. The observed anti-cancerous and the anti-microbial activities of the MOSs extracts can be attributed to the charisma of various bioactive compounds including the oleic acid, palmitic acid, hexadecenoic acid ethyl ester and D-allose.

**Conclusion:** Current observations may contribute towards the development of the MOSs-based biomedicine (organic without any side effects, cheap, plentiful, pure and sustainable) effective for the cancer and bacterial infection cure.

## 1. Introduction

Generally, the moringa tree with the scientific name of Moringa oleifera Lam. (MOL) is the part of Moringaceae family [1]. Being originated from the Northern India and Pakistan region alongside the Himalaya Mountains, moringa tree also grows in different parts of African, Asian, and South American tropics [2-4]. In Pakistan, India, Thailand and Philippines the moringa leaves and fruits are usually consumed as the vegetables [5]. In recent times, several countries (Mexico, Hawaii, Cambodia, and the Caribbean Islands) started cultivating it due to its numerous health benefits and nutritional values [6,7]. Apart from the staple organic food with enriched vitamins, proteins and minerals, the natural moringa leaves and fruits have several biological applications [5,8]. The MO as a food is recommended by the WHO due to its ramified nutritionals benefits to human health [5].

In the past, several emperors worldwide used to consume the fruits and leaves of the Moringa plants for keeping the skin healthy, acquiring energy, relieving the tension/stress and pain during the battles [8]. Intensive studies revealed that the MO have many notable attributes including anti-oxidant, anti-microbial, anti-cancer, anti-inflammatory, anti-ulcer, anti-hypertensive, anti-urolithiatic, anti-diabetic, anti-asthmatic, anti-aging, cardiovascular, analgesic, immunomodulation, hepatoprotective, diuretic, anthelmintic, and hypoglycemic characteristics [2,9-19]. In fact, the presence of some distinctive chemical compounds and elements such as the 4-(4′-O-acetyl-α-L-rhamnopyranosyloxy) benzyl isothiocyanate, niazimicin, 4-(α-L-rhamnopyranosyloxy) benzyl isothiocyanate, pterygospermin, 4-(α-Lrhamnopyranosyloxy), benzyl isothiocyanate, and benzyl glucosinolate impart the hypo-tensive, anti-bacterial and anti-cancer efficacies to the MO leaves and fruits [20]

To take the advantages of the MO plant products as the food supplements and medicines as well as to determine its other health benefits various techniques have been developed for extracting the contents from the MO leaves and fruits. These methods include pressurized liquid extraction [21], ultrasound [22], microwave-based extraction [23], and supercritical fluid extraction [24]. However, all these techniques suffer from some limitations in terms of the requirement of a large quantity of toxic organic solvents, cumbersome sample preparation procedures and costly [25]. To overcome such drawbacks the LIBS has been emerged as an efficient approach to determine the elemental compositions (various trace elements) in the extracts obtained from different medicinal plants.

Currently, the laser induced breakdown spectroscopy (LIBS) technique has widely been exploited for elemental analyses of different types of samples. It is a versatile analytical technique with many distinct features such as the real-time measurement, rapidity, cost-effectiveness, sensitiveness, and in-situ elemental analysis with microanalyses capacity for diverse types of materials and phases (solid, liquid, or gas) [26-31]. In addition, this technique is simple, eco-friendly, and does not need cumbersome sample preparation protocol because the sample treatments frequently tend to induce errors through contamination and losses. Depending on the varieties of the inorganic constituents present in the medicinal plants, the LIBS technique can accurately disclose their secrecy in a scientific manner. It is needless to mention that, besides the occurrences of diverse organic molecular components most of the edible plants (herbs and vines) are enriched with inorganic molecules and elements (in the form of vitamins, proteins and minerals), signifying their immense health benefits. Since ancient times, the Ayurvedic medicinal practices have been relied on these crude chemical components extracted from the medicinal plants [32,33]. The screening of the glycemic index of various elements in diverse medicinal plants was performed to evaluate their anti-diabetic potency [34,35].

Based on the abovementioned factors, for the first time we used the LIBS technique to analyze the presence of various trace elements in the MO seeds (MOSs) extracts. The gas chromatography - mass spectroscopy (GC-MS) measurement of the ethanolic MOSs extract was carried out to validate the LIBS results. In addition, the anti-cancerous and anti-microbial characteristics of the MOSs extracts were evaluated. The human colorectal carcinoma (HCT-116) cancerous cells and embryonic kidney (HEK-293) normal cells were chosen for the anti-cancerous assessment (using MTT assays) of the MOSs extracts. The *gram-positive Staphylococcus aureus* (*S. aureus*) and *Escherichia coli* (*E. coli*) bacterial strains were selected for the bactericidal evaluation (using Agar well diffusion) of the MOSs extracts. Results were interpreted and discussed to understand the mechanism of the anti-cancerous and anti-microbial effectiveness of such extracts.

## 2. Experimental procedure

### 2.1. Seeds collection and extract synthesis

The good quality MOSs were purchased from the local market (originally imported from India) and used for the extract preparation in few steps. The coatings of the seeds were first removed before being crushed and ground to fine powder. Next, the resultant powder was compressed in the form of pellets. Fig.1 (a)-(d) shows various stages of the sample preparation for further analyses using the LIBS technique. Various amounts of aliquots (5, 10, 15, 20 and 25 g) were mixed separately in 200 ml of deionized water (DIW). The obtained solution was heated at 60 °C with constant stirring and then kept overnight before being filtered (Whatman filter paper No.1, Sigma-Aldrich). The obtained filtrate was maintained at 4 °C for the anti-cancerous activity analyses of the MOSs. An alcoholic extraction was prepared for the antimicrobial activity test. In this process, the fine powder of MOSs of various contents (5, 10, 15, 20 and 25 gm) was mixed in ethanol (200 ml) separately and stirred for 5 h. Later, the solutions containing the MOSs were filtered and placed inside a rotary evaporator.

**Fig 1.**
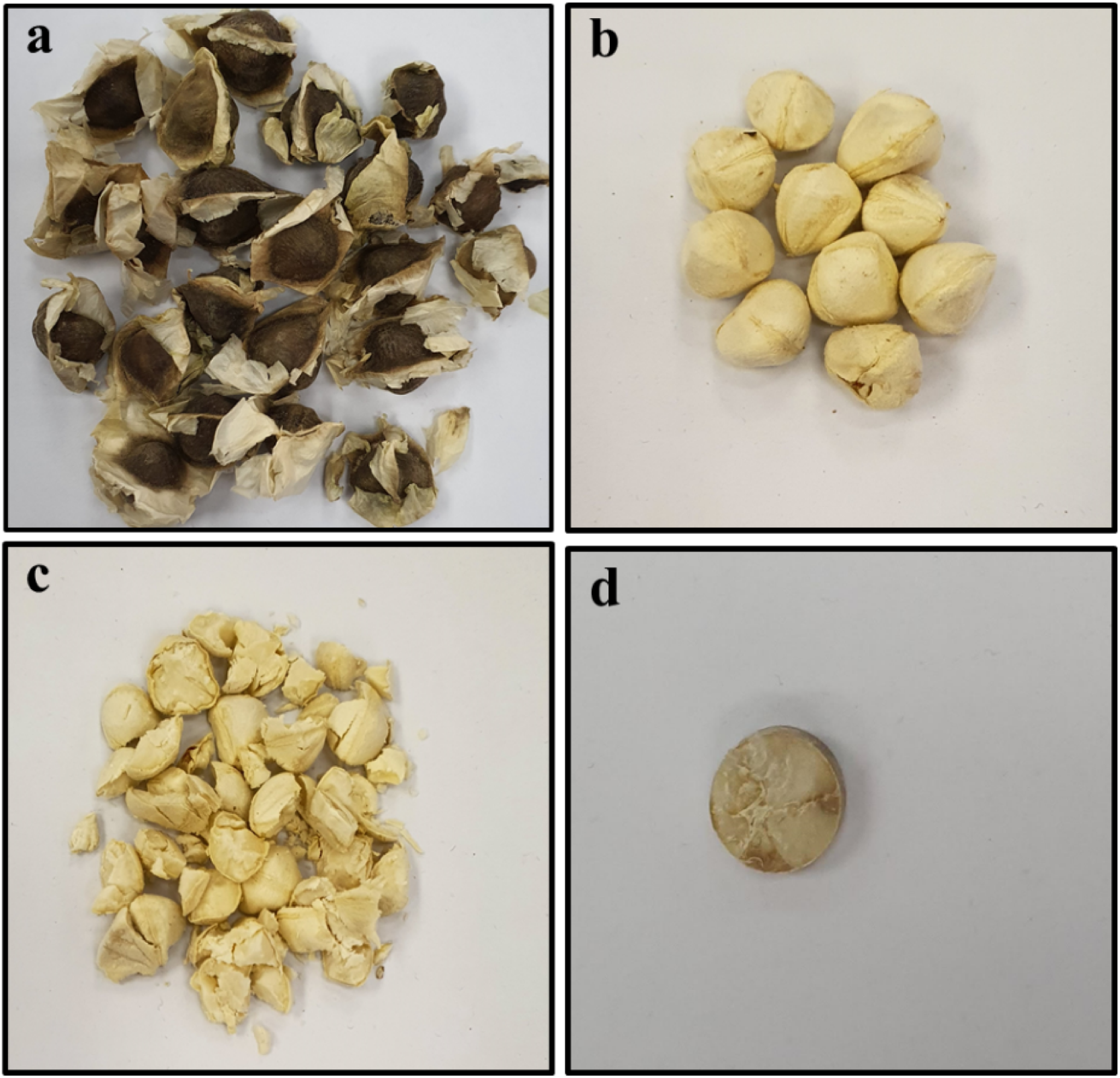
*Moringa oleifera* seeds sample: **a)** seed, **b)** seeds without coat, **c)** crushed seeds, **d)** seeds in the form of pellets.

### 2.2. LIBS Setup Details

Fig 2 depicts the customized LIBS setup used to quantify the constituent elements present in the MOSs samples. It consisted of quadrupled Q-switched Nd:YAG laser (model QUV-266-5) operated at the wavelength of 266 nm, repetition rate of 20 Hz, maximum output energy of 30 mJ and pulse width of 8 ns. The collimated laser pulses were focused onto the MOSs pellet (target) using a UV convex lens of 30 mm focal length for ablation. Upon ablating the target pellets a plasma plume was generated. The emitted plasma was detected/collected using the fiber optical system equipped with a miniaturized lens placed at an angle of 45°. The other end of fiber was connected to the 500 mm spectrograph (Andor SR 500i-A) with the grating groove of approximately 1200 lines/mm. The pellet was mounted on a 2-D motorized sample holder capable of moving in the X-Y direction to avoid the crust formation on the target surface due to multiple laser shots on the same spot. The emission spectra were recorded by a ICCD camera (model iStar 320T, 690 × 255 pixels with delay time setting at 300 ns) and transferred to the interfaced on line personal computer (PC).

**Fig. 2.**
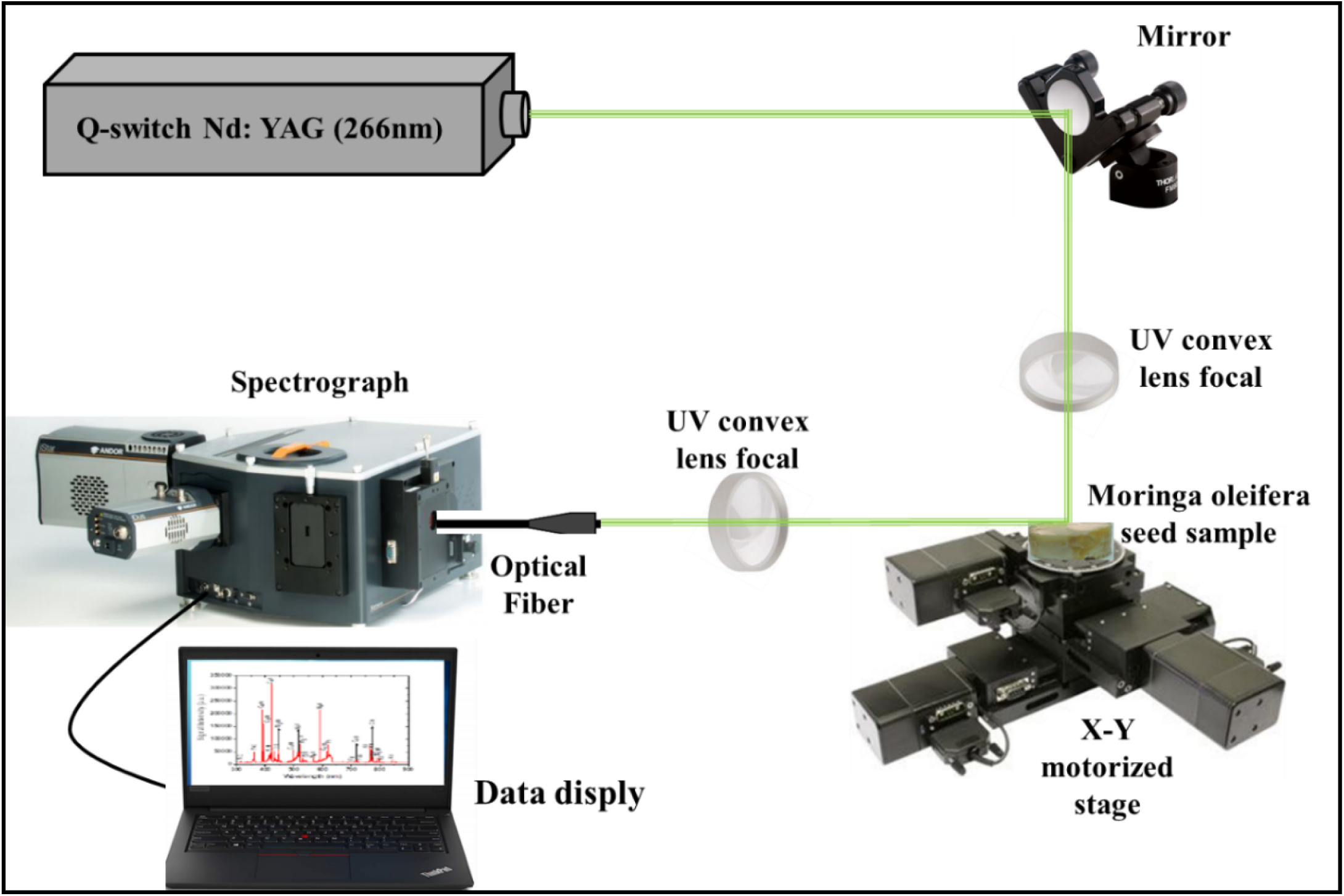
Schematic of our LIBS setup, where the inset shows an image of *Moringa oleifera* seed sample.

### 2.3 GC-MS measurement

The ethanolic MOSs extract was analyzed using GC-MS (Shimadzu GC-2010 Plus) which was equipped with a split/spitless auto-injector (AOC-20i series) and coupled to a QP2010 Ultra single quadrupole (Shimadzu Corporation, Kyoto, Japan). The GC separation was achieved via an Rxi-5MS fused silica capillary column (Restek, USA) of dimension (30 m × 0.25 mm id × 100 µm film thickness). The temperature was raised from 60 °C (kept for 0.5 min) to 280 °C at the rate of 5 °C/min (hold for 5 min). The inlet was operated in the splitless mode at the temperature of 270 °C. Helium was flown (purity of 99.9999%) as the carrier gas at the rate of 1 ml/min and the temperature of the MS transfer line was maintained at 280 °C. The ion source was worked in the electron impact mode at the energy of 70 eV and temperature of 250 °C. The full scan mass spectra were collected from 33 to 550 m/z. The spectral data was obtained by controlling the GC-MS and processed using the GC-MS solution (version 4.52, Shimadzu Corporation, Japan). The detected volatile compounds were identified using the NIST 11 and WILEY 9 libraries and relative area of each compound was calculated.

### 2.4. Anti-cancerous activity evaluation of MOSs extract

#### 2.4.1. In vitro cell culture and cell viability test

In this study, the cancerous and normal (healthy) cell line including the respective human colorectal carcinoma (HCT-116) and embryonic kidney (HEK-293) cells were used to evaluate the anti-cancerous activity of various MOSs extracts on these cells. Following the earlier prescribed method [36,37], the proposed cells were seeded in 96 well plates and grown in the Dulbecco’s Modified Eagle Medium (DMEM) supplemented with the reagents such as selenium chloride, L-glutamine, fetal bovine serum, antibiotic penicillin and streptomycin. One group of these cells were cultured inside a CO_2_ incubator at the temperature of 37 °C and treated at various concentrations (0.030 mg/ml to 0.1 mg/ml) of the MOSs extracts. Another group was treated at different concentrations (0.030 mg/ml to 0.1 mg/ml) of the MOSs extracts. The control group was devoid of the MOSs extracts. After treating the cells using the MOSs extracts for 48 hrs they were treated with 3-[4,5-dimethylthiazol-2-yl]-2,5-diphenyl-tetrazolium bromide (MTT assay, Molecules, New Zealand) for 4 hrs. Next, the growth medium was eliminated from the plates and dimethyl sulfoxide (DMSO) was incorporated in every well where MTT created the Formazan crystals. Later, the culture plates were inspected at the wavelength of 570 nm using a microplate reader (Bio-Rad Laboratories, Hercules, CA, USA). Finally, the GraphPad Prism Software was used to analyze the acquired data where a one-way analysis of variance (ANOVA) statistical tool was utilized.

#### 2.4.2. Nuclear staining via DAPI

The cancer cells (HCT-116) were cultured inside the CO_2_ incubator at 37 °C and treated for 48 hrs using the MOSs extract of concentration 0.066 mg/ml. The control group was made without adding any such extract. The effects of the MOSs extracts on the cells nuclei were studied after staining by DAPI. Next, the cold paraformaldehyde (4%) was used to pre-treat these cells and washed using Triton X-100 (0.1%) prepared in the phosphate buffer saline (PBS). Then, both control cell and those treated with the MOSs extracts were stained using DAPI (1.0 μg/ml) made in the PBS. Lastly, the cells were rinsed using Triton X-100 (0.1%) made in PBS. The morphologies of the control cells and those treated by MOSs extracts were scanned via a confocal scanning microscope (CSM, Zeiss, Germany).

### 2.5. Anti-microbial activity assessment of MOSs extract

The bactericidal efficacy of the proposed MOSs extracts at various concentrations (50, 100, 150, 200, 250 mg/ml) was assessed using the Agar well diffusion method. The *gram-positive S. aureus* and *E. coli* bacterial strain were selected. For the preparation of inoculum, bacteria were freshly grown in the nutrient broth (NB) at 37 °C by incubating overnight and regulated to the 0.5 McFarland standards. The Mueller Hinton agar solution was prepared by adding 100 µl of freshly adjusted inoculum. The inoculum was spread uniformly over the surface of the MHA plates and left to dry under the aseptic conditions. Consequently, the sterile cork-borer was used to punch the inoculated plates for the wells of 6 mm. Approximately 50 μl of the prepared extract was placed in the wells at 37 °C for overnight incubation. At the end of the incubation period, the bacterial zone of inhibition (ZOI) diameter around the wells was measured to evaluate the inhibition effectiveness of the studied MOSs extracts [38].

## 3. Results and Discussion

### 3.1. Qualitative analysis of MOSs using LIBS

The LIBS spectra (Fig 3) of the MOSs were recorded in range of 200 to 800 nm. For detecting the major elements, the spectra were collected from different regions of the pellets each of scan length 50 nm. In order to reduce the background noise that may affect the quality of the LIBS signal, the LIBS parameters including the delay times, number of accumulations and laser energies were optimized. The recorded LIBS signal of the MOSs comprised of various significant spectral peaks of the atomic and ionic lines (with varying intensities) were related to abundant elements. Based on the NIST database the observed spectral lines were identified and categorized in terms of the characteristic elements present in the MOSs, suggesting their significant role towards the anti-microbial and anti-cancerous activities. Clearly, the measured LIBS spectra (Fig 3) showed the presence of Ca, K, Mg, P, S, Fe, Mn, Zn, Na and Se.

**Fig 3.**
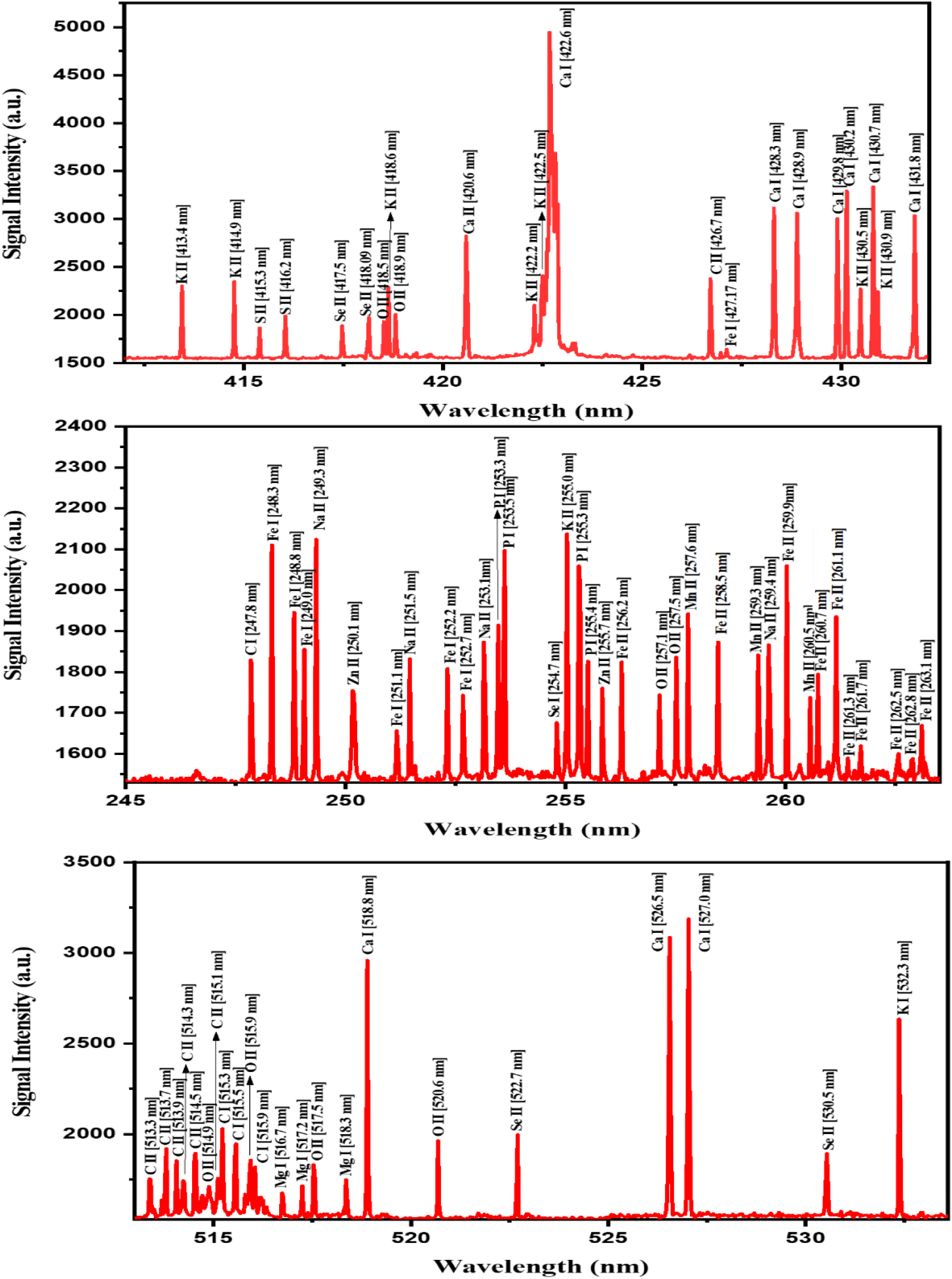
LIBS spectra of *Moringa oleifera* seeds.

Table 1 presents the LIBS signal intensities of the spectral transition lines corresponding to various detected elements in MOSs. In accordance to the Boltzmann distribution, the intensities of the LIBS spectral lines have direct relationship with elemental contents in MOSs [39]. This correlation was attained by considering the intensity ratio of the detected elemental lines with that of the C line taken as the reference (247.8 nm). The achieved intensity ratio of the elements Ca, K, Mg, P, S, Fe, Mn, Zn, Na and Se were ranged from 2.7 - 1.7, 1.2 - 1.4, 0.9, 1.1, 1.1, 1.1 - 0.9, 1.0, 0.9, 1.2 and 1.1, respectively which were consistent with those reported in the literature [40]. These results approved that the MOSs are rich in different minerals which are useful to human as food and medicine, indicating their remarkable influence in regulating the level of blood pressure, blood lipids, regulating the stomach, protecting the liver, strengthening bones, generate protein and enhancing the immunity of the body [41]. Furthermore, the existence of Se in the MOSs seeds plays a vital role by protecting from cancer, cardiovascular disease, cognitive decline and thyroid disease. Moreover, these seeds exhibit a powerful antibacterial activity due to the availability of the detected elements including Ca, K, Mg, P, S, Fe, Mn, Zn, Na and Se [42].

**Table 1.**
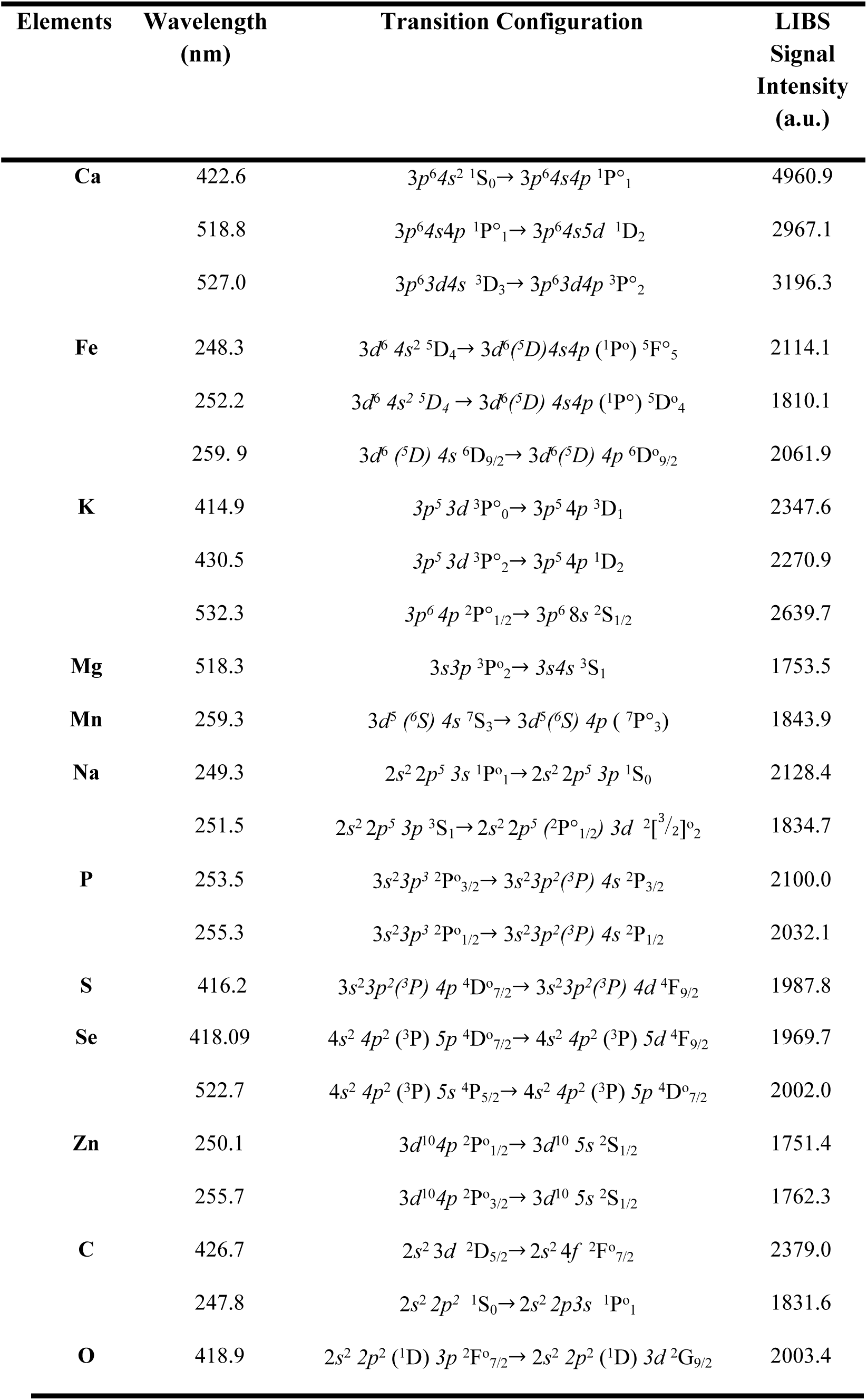
Some of the detected spectral lines of the elemental analytes present in the Moringa seeds recorded by using our LIBS system.

### 3.2. Volatile content analyses of MOSs using GC-MS

A total of 114 volatile compounds were detected in the proposed MOSs via GC-MS data analyses. These compounds included different chemical groups including varieties of fatty acid, aldehyde, ester, alcohol, ketone and hydrocarbon. Table 2 and Fig 4 show the retention times and percentage composition of all the identified chemical compounds existed in the MOSs. The detected major constituents were the oleic acid (22.53%), 2,3-dihydroxypropyl elaidate (13.48%), 9-octadecenoic acid (Z)-, 2,3-dihydroxypropyl ester (11.35%), docosenamide (6.04%), ethyl oleate (6.03%), 1,3-propanediol, 2-ethyl-2-(hydroxymethyl) (5.52%), oleic anhydride (3.96%) and 2-Propanone, 1,1-dimethoxy (3.86%). These MOSs are rich in the fatty acids and their ester derivatives (65.45%) followed by alcohols (9.4%), nitrogen containing compounds (9.09%), ketones (5.34%) and aldehydes (2.88%). The fatty acids and alcohols in plants may undergo esterification to form esters [43]. The GC-MS measurement revealed the occurrences of several fatty acids and their esters such as oleic, n-hexadecenoic (palmitic), cis-9-Hexadecenoic acid (Palmitoleic), octadecanoic (stearic) acids and their alcohols and esters. In fact, most of such compounds have been reported to possess anti-cancer activity. For example, the D-allose was reported to inhibit cancer cells growth at G1 phase [44]. The palmitic acid disclosed a selective cytotoxicity against the leukemic cells in humans [45]. In addition, such fatty acids are known to have antifungal and antibacterial effects [46]. The oleic acid that was identified as the major compound in the MOSs is an omega-9-fatty acid with numerous benefits for the human health. This can effectively be used to prevent the ulcerative colitis [47] and reduce the blood pressure [48]. On top, it has remarkable antioxidant efficiency [49].

**Table 2:** Volatile compounds identified in Moringa *oleifera* seeds using GC-MS analysis

| No. | Compounds | RT | Peak area (%) | No. on Chromatogram |
| --- | --- | --- | --- | --- |
| **Esters** |  |  |  |  |
| 1 | Propanoic acid, 2-oxo-, methyl ester | 6.29 | 0.76 | 3 |
| 2 | Acetic acid, ethoxyhydroxy-, ethyl ester | 7.04 | 0.04 | 5 |
| 3 | Diethoxymethyl acetate | 12.83 | 0.09 | 28 |
| 4 | 2-Propenoic acid, 2-methyl-, 2-hydroxypropyl ester | 13.13 | 0.02 | 29 |
| 5 | 6,9,12-Octadecatrienoic acid, phenylmethyl ester, (Z,Z,Z)- | 17.80 | 0.01 | 42 |
| 6 | Cyclopentanecarboxylic acid, 4-tridecyl ester | 19.18 | 0.03 | 47 |
| 7 | 1-Cyclohexene-1-carboxylic acid, 2,6,6-trimethyl-, methyl ester | 20.06 | 0.13 | 49 |
| 8 | Hexanoic acid, 4-hexadecyl ester | 28.29 | 0.07 | 64 |
| 9 | Phthalic acid, diethyl ester | 29.57 | 0.14 | 68 |
| 10 | 2-Propenoic acid, pentadecyl ester | 31.63 | 0.59 | 70 |
| 11 | Acetic acid, 3,7,11,15-tetramethyl-hexadecyl ester | 33.63 | 0.02 | 72 |
| 12 | Phthalic acid, dibutyl ester | 35.69 | 0.07 | 77 |
| 13 | Phthalic acid, diisobutyl ester | 35.76 | 0.15 | 78 |
| 14 | Palmitoleic acid, methyl ester | 36.29 | 0.03 | 79 |
| 15 | Pentadecanoic acid, 13-methyl-, methyl ester | 36.64 | 0.09 | 80 |
| 16 | 9-Hexadecenoic acid, ethyl ester | 37.65 | 0.15 | 86 |
| 17 | Hexadecanoic acid, ethyl ester (Ethyl palmitate) | 37.99 | 0.78 | 88 |
| 18 | Heptadecanoic acid, ethyl ester | 38.42 | 0.46 | 90 |
| 19 | Propanoic acid, 3-mercapto-, dodecyl ester | 38.59 | 0.25 | 91 |
| 20 | 9-Octadecenoic acid, methyl ester, (E)- (Methyl elaidate) | 40.16 | 1.52 | 94 |
| 21 | Oleic acid, methyl ester (Methyl Oleate) | 40.26 | 0.15 | 95 |
| 22 | l-(+)-Ascorbic acid 2,6-dihexadecanoate | 40.56 | 0.10 | 96 |
| 23 | Ethyl Oleate | 41.39 | 6.03 | 98 |
| 24 | Octadecanoic acid, ethyl ester (Ethyl stearate) | 41.77 | 0.53 | 100 |
| 25 | 9-octadecenyl ester (Oleyl oleate) | 43.52 | 0.34 | 102 |
| 26 | 2,3-dihydroxypropyl elaidate | 43.79 | 13.48 | 104 |
| 27 | 9-Octadecenoic acid, 1,2,3-propanetriyl ester | 44.29 | 0.90 | 105 |
| 28 | Docosanoic acid, ethyl ester | 45.27 | 0.37 | 107 |
| 29 | Oleoyl chloride | 46.07 | 0.65 | 109 |
| 30 | 9-Octadecenoic acid (Z)-, 2,3-dihydroxypropyl ester | 47.10 | 11.35 | 111 |
| 31 | Glycidol stearate | 47.52 | 1.76 | 113 |
| 32 | Hexadecanoic acid, 2-hydroxy-1-(hydroxymethyl)ethyl ester | 47.80 | 1.57 | 114 |
| **Alcohols** |  |  |  |  |
| 33 | 1,4-Cyclohexanediol, trans- | 7.22 | 0.02 | 6 |
| 34 | 1,2-Propanediol, 3-methoxy- | 7.43 | 0.05 | 7 |
| 35 | Ethanol, 2,2-diethoxy- | 7.54 | 0.31 | 8 |
| 36 | 1,2,4-Butanetriol | 9.50 | 0.04 | 15 |
| 37 | Glycerin | 11.18 | 1.35 | 21 |
| 38 | 1-Butanol, 4-(ethylthio)- | 11.74 | 0.02 | 22 |
| 39 | Methoxyacetaldehyde diethyl acetal | 11.88 | 0.02 | 24 |
| 40 | 1-Dodecanol | 18.43 | 0.48 | 43 |
| 41 | 1-Tetradecanol | 24.15 | 0.42 | 58 |
| 42 | 1,3-Propanediol, 2-ethyl-2-(hydroxymethyl)- | 25.44 | 5.52 | 60 |
| 43 | 1-Tridecanol | 26.33 | 1.06 | 61 |
| 44 | 3-Hexadecanol | 31.35 | 0.05 | 69 |
| 45 | 3-Heptadecanol | 35.56 | 0.06 | 76 |
| **Aldehydes** |  |  |  |  |
| 46 | Butanal, 3-hydroxy- | 8.38 | 0.00 | 12 |
| 47 | Heptanal | 10.01 | 0.02 | 17 |
| 48 | Octanal | 12.57 | 0.04 | 27 |
| 49 | Nonanal | 15.79 | 0.09 | 36 |
| 50 | 2-Methyl-oct-2-enedial | 20.61 | 0.04 | 51 |
| 51 | Undecanal | 23.52 | 0.02 | 54 |
| 52 | Tridecanal | 34.46 | 0.01 | 74 |
| 53 | 10-Octadecenal | 37.42 | 0.09 | 84 |
| 54 | 9-Octadecenamide | 37.55 | 0.09 | 85 |
| 55 | cis-9-Hexadecenal | 38.24 | 0.55 | 89 |
| 56 | cis-13-Octadecenal | 47.23 | 1.92 | 112 |
| **Ketones** |  |  |  |  |
| 57 | 2-Propanone, 1,1-dimethoxy- | 6.84 | 3.86 | 4 |
| 58 | 2-Butanone | 8.15 | 0.03 | 10 |
| 59 | Dihydroxyacetone | 8.97 | 0.14 | 13 |
| 60 | 1,2-Cyclopentanedione | 10.17 | 0.14 | 18 |
| 61 | 2-Propanone, 1-(1,3-dioxolan-2-yl)- | 10.87 | 0.04 | 20 |
| 62 | 1,3-Dioxol-2-one,4,5-dimethyl- | 13.53 | 0.28 | 30 |
| 63 | 2-Heptanol, 5-ethyl- | 14.31 | 0.01 | 33 |
| 64 | 2-Methyl-4-octanone | 16.47 | 0.03 | 37 |
| 65 | 2-Pentanone, 3,4-epoxy- | 18.93 | 0.03 | 45 |
| 66 | 1-Oxa-spiro[4.5]deca-6,9-diene-2,8-dione | 37.01 | 0.05 | 82 |
| 67 | Z-11-Pentadecenol | 39.21 | 0.05 | 92 |
| 68 | Cyclopentadecanone, 2-hydroxy- | 39.82 | 0.13 | 93 |
| 69 | Cyclopentadecanone | 43.43 | 0.21 | 101 |
| 70 | 2-Tetradecanone | 46.69 | 0.34 | 110 |
| **Acids** |  |  |  |  |
| 71 | Acetic acid, (acetyloxy)- | 5.88 | 0.18 | 2 |
| 72 | Butanoic acid, 3-hydroxy- | 10.59 | 0.03 | 19 |
| 73 | Octanoic acid | 17.42 | 0.03 | 41 |
| 74 | Nonanoic acid | 20.36 | 0.02 | 50 |
| 75 | n-Hexadecanoic acid (Palmitic acid) | 37.31 | 0.05 | 83 |
| 76 | Oleic Acid | 40.94 | 22.53 | 97 |
| **Furans and lactones** |  |  |  |  |
| 77 | Furfural | 7.62 | 0.02 | 9 |
| 78 | 2(5H)-Furanone | 9.84 | 0.04 | 16 |
| 79 | 2-Hydroxy-gamma-butyrolactone | 12.08 | 0.23 | 25 |
| 80 | 2,5-Dimethyl-4-hydroxy-3(2H)-furanone | 14.10 | 0.19 | 32 |
| 81 | 1,2-Ethanediol, 1-(2-furanyl) | 19.10 | 0.03 | 46 |
| 82 | 5-Hydroxymethylfurfural | 19.45 | 0.36 | 48 |
| 83 | 3-Deoxy-d-mannoic lactone | 28.92 | 0.52 | 65 |
| **Nitrogen containing compounds** |  |  |  |  |
| 84 | N,N-Dimethylaminoethanol | 5.22 | 0.12 | 1 |
| 85 | 1,3,5-Triazine-2,4,6-triamine | 14.78 | 0.68 | 34 |
| 86 | Acetic acid, 2-(N-methyl-N-phosphonomethyl)amino- | 15.62 | 0.11 | 35 |
| 87 | 1-Heptadecanamine | 18.67 | 0.07 | 44 |
| 88 | Nonanamide | 23.03 | 0.02 | 52 |
| 89 | Dodecanamide (Lauryl amide) | 33.33 | 0.04 | 71 |
| 90 | Tetradecanamide (Myristic amide) | 37.72 | 0.44 | 87 |
| 91 | Hexadecanamide (Palmitic amide) | 41.72 | 1.26 | 99 |
| 92 | Docosenamide | 44.99 | 6.04 | 106 |
| 93 | Nonadecanamide | 45.35 | 0.32 | 108 |
|  | ***Sulfur containing compounds*** |  |  |  |
| 94 | Sulfurous acid, cyclohexylmethyl hexadecyl ester | 23.17 | 0.67 | 53 |
|  | ***Hydrocarbons*** |  |  |  |
| 95 | 1-Butene, 4,4-diethoxy-2-methyl- | 9.43 | 0.07 | 14 |
| 96 | 1-Methyl-2-octylcyclopropane | 12.17 | 0.07 | 26 |
| 97 | 2-Trifluoroacetoxytridecane | 13.92 | 0.03 | 31 |
| 98 | trans-2,3-Epoxy-nonane | 23.64 | 0.03 | 55 |
| 99 | 2-Heptafluorobutyropentadecane | 23.89 | 0.04 | 56 |
| 100 | 1,2-Epoxyundecane | 24.02 | 0.02 | 57 |
| 101 | Heptacosane | 24.36 | 0.03 | 59 |
| 102 | 1-Heptadecene | 29.26 | 0.33 | 66 |
| 103 | Octadecane, 1,1'-[(1-methyl-1,2-ethanediyl)bis(oxy)]bis- | 29.43 | 0.01 | 67 |
| 104 | 1-Nonadecene | 33.85 | 0.22 | 73 |
|  | ***Pyrans*** |  |  |  |
| 105 | Tetrahydro-4H-pyran-4-ol | 17.07 | 0.03 | 38 |
| 106 | 4H-Pyran-4-one, 2,3-dihydro-3,5-dihydroxy-6-methyl- | 17.20 | 0.02 | 39 |
|  | ***Others*** |  |  |  |
| 107 | 5,6-Dihydroxypiperazine-2,3-dione dioxime | 8.32 | 0.02 | 11 |
| 108 | Tetraethyl silicate | 11.81 | 0.04 | 23 |
| 109 | Silanediol, dimethyl-, diacetate | 17.30 | 0.07 | 40 |
| 110 | D-Allose | 26.61 | 0.19 | 62 |
| 111 | Phenol, 2,4-bis(1,1-dimethylethyl)- | 27.43 | 0.51 | 63 |
| 112 | Oxirane, hexadecyl- | 34.90 | 0.02 | 75 |
| 113 | Ethyl iso-allocholate | 36.94 | 0.06 | 81 |
| 114 | Oleic anhydride | 43.70 | 3.96 | 103 |

\* RT: Retention time in minutes.

**Fig 4.**
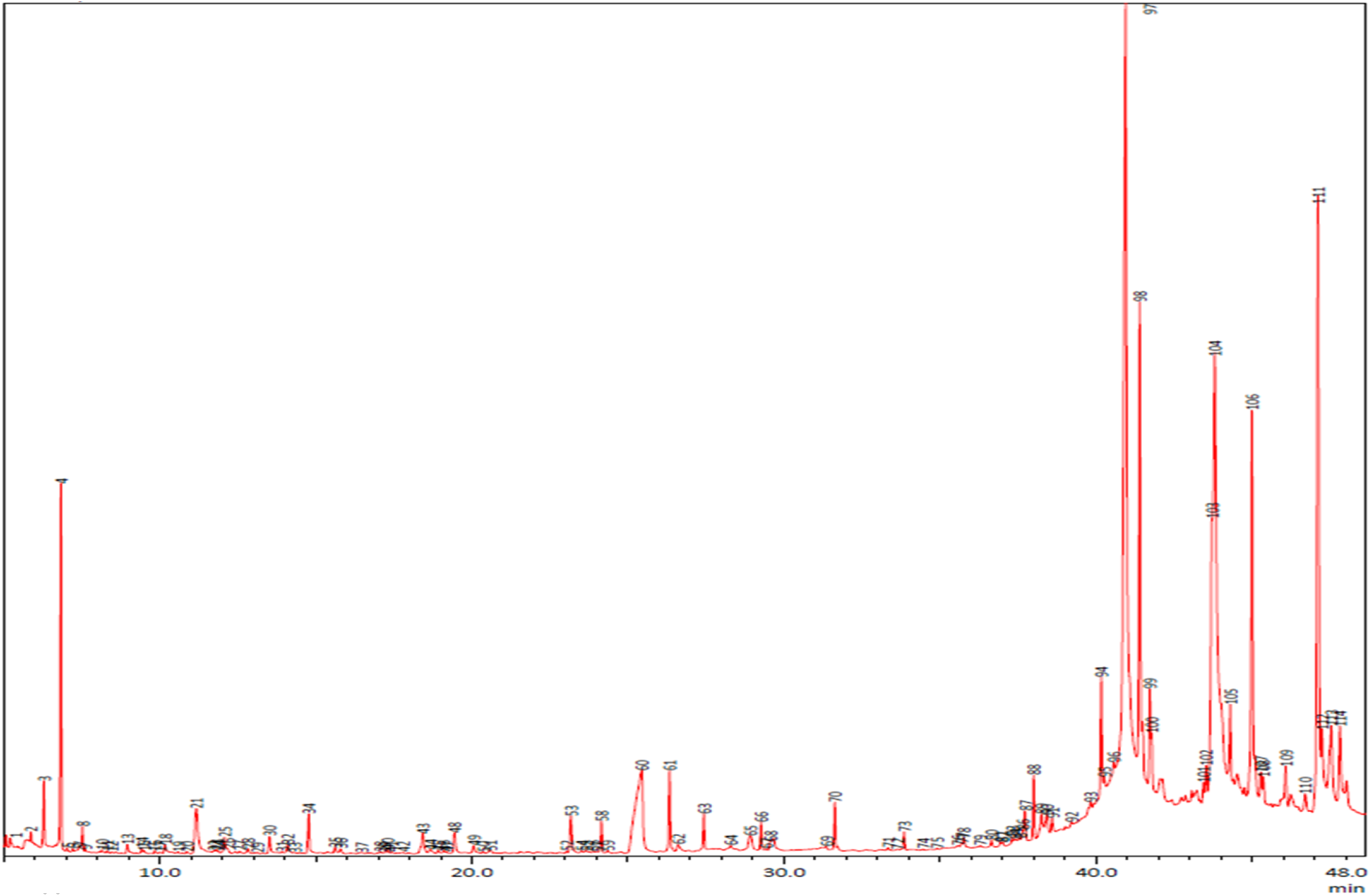
GC-MS chromatograph of Moringa *oleifera* seeds.

### 3.3. Anti-proliferative activities of MOSs

The anti-proliferative effectiveness of the MOSs extracts were examined on the colon cancer cells via the MTT assay. After 48 hrs of treatments, the cell viability status was examined in both normal (HEK-293) and cancerous (HCT-116) cells. The percentage of cell viability and inhibitory concentration (IC_50_) values of both normal and treated cells were calculated. The MTT assay results after 48 hrs of treatment revealed the cytotoxic effects of the MOSs extracts, indicating their inhibitory action on the HCT-116 (cancerous) cells. The average of cell viability data showed (Fig 5) that the treatment using MOSs extracts could induce a significant decrease in the cancer cells viability compared to the control cells (without treatment using MOSs extracts).

**Fig 5.**
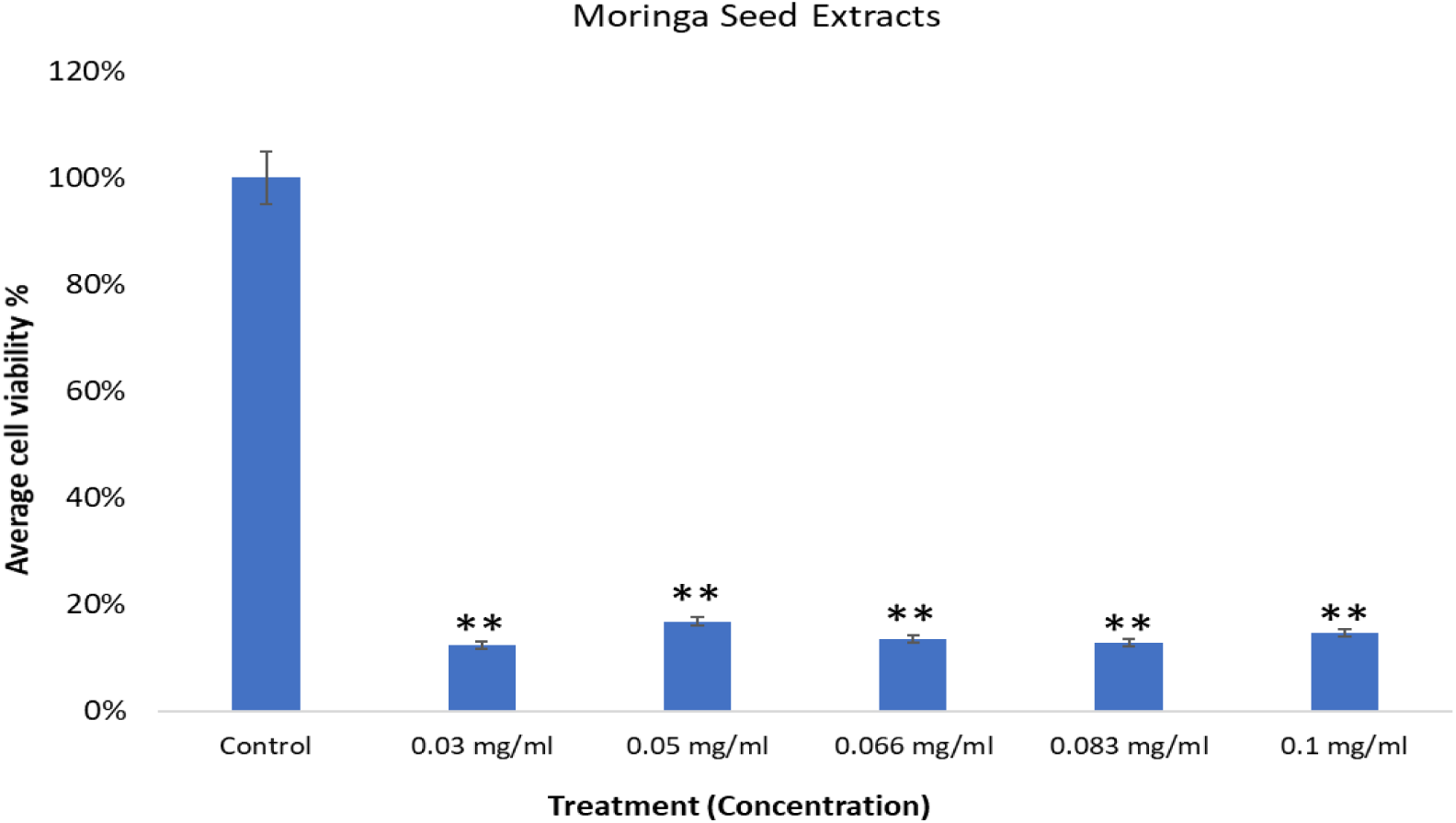
Impact of *Moringa oleifera* seed extracts on the colon cancer cells. The HCT-116 treated with moringa seed extracts for 48 hrs with different concentrations showing the average cell viability done by MTT assay. (**p < 0.01)

The specificity of the MOSs extracts (at different concentrations ranged from 0.03 to 0.1 mg/ml) on the normal and healthy (HEK-293) cells were inspected after 48 hrs of treatment using the MTT assays (Fig 6). The results showed that the inhibitory activity of the MOSs extracts (irrespective of their concentration) on the normal cells was insignificant, thereby suggesting the specificity of the MOSs extract towards the colon cancer cells only.

**Fig 6.**
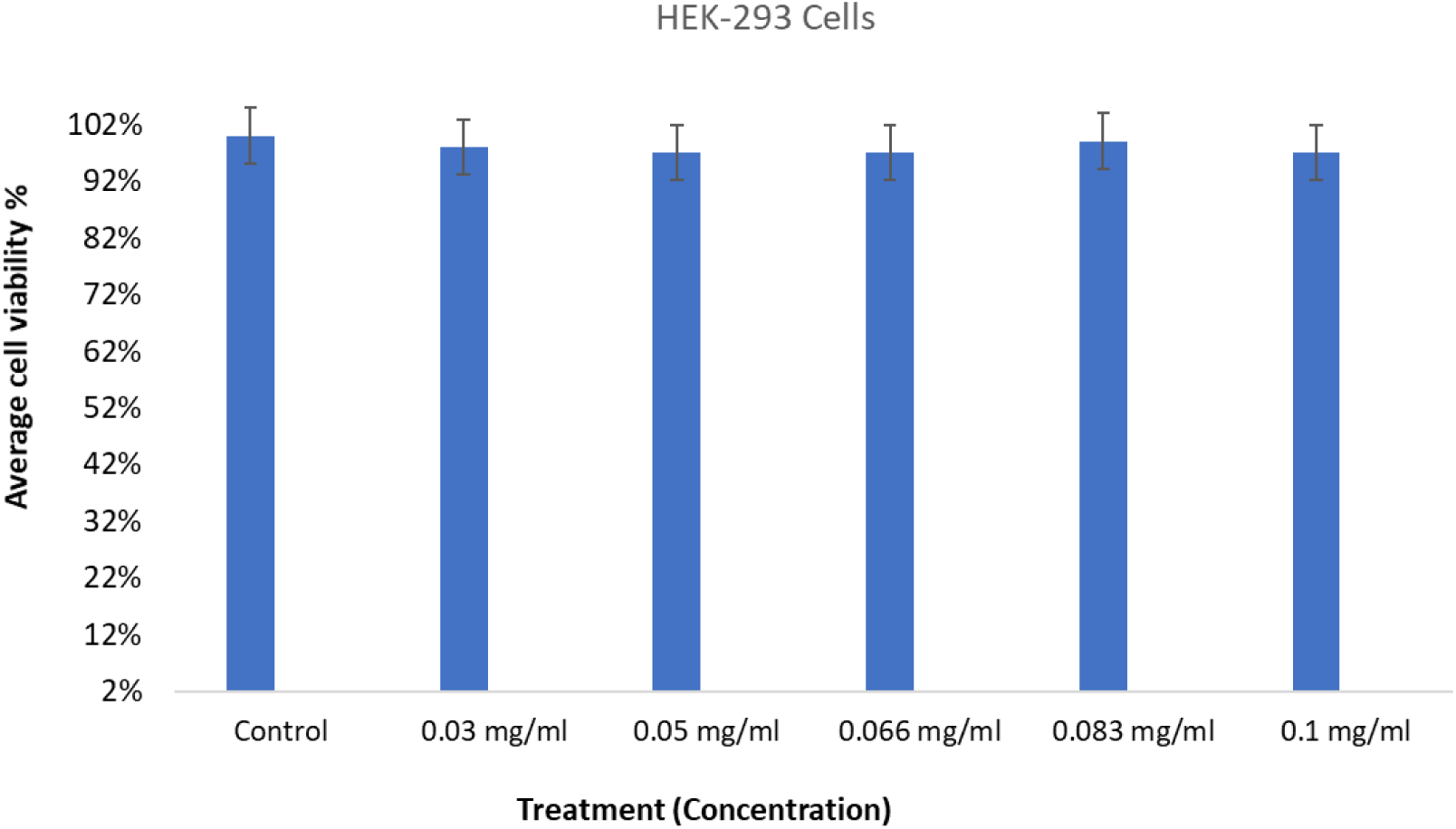
Impact of *Moringa oleifera* seed extracts on HEK-293 which were treated with moringa seed extracts for 48h showing the average cell viability done by MTT assay. The cells were treated with different concentrations (0.03 to 0.1 mg/ml) and. Data are the means ± SD of three different experiments. Difference between two treatment groups were analysed by student’s t test

### 3.4. Nuclear disintegration due to treatment of Moringa seed extracts

The morphologies of the untreated and MOSs extracts treated cells were imaged using the CSM as shown in Fig 7. The cancerous cells (HCT-116) treated with MOSs extracts exhibited strong inhibitory action (Fig 7(b)) than the control sample (Fig 7(a)).

**Fig 7.**
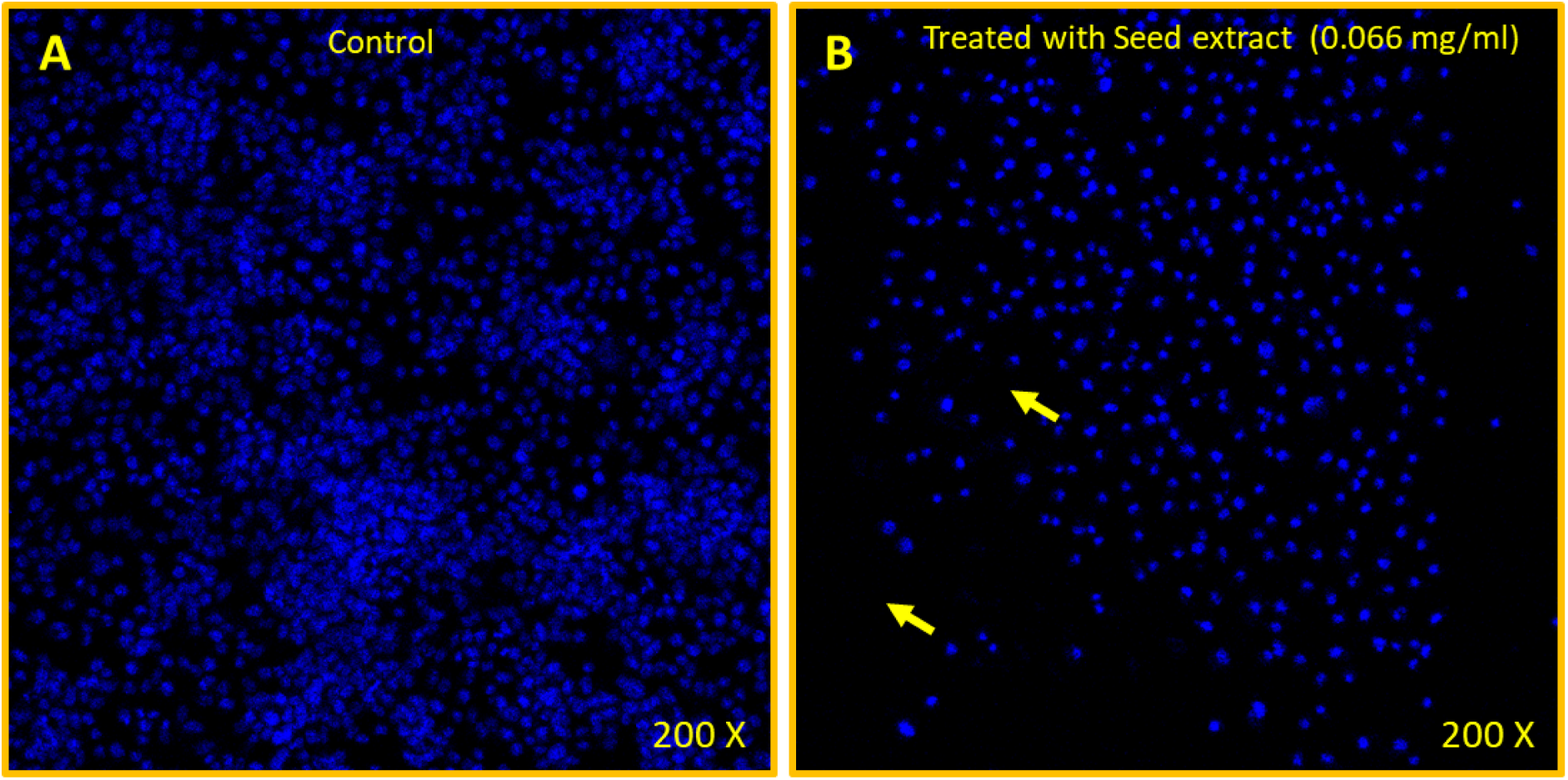
Nuclear staining by DAPI. The cancerous cells HCT-116 cells. (A) Control (nontreated) (B) treated with seed extracts of *Moringa oleifera* (0.066 mg/mL). DAPI stained cells visualized through confocal microscope and figure B show significant loss (arrows) of staining due to treatment. 200x magnifications.

It was inferred that the observed strong anti-cancerous activities of the MOSs extracts on the colon cancer cells may be due to the presence of high level of oleic acid and fatty acid in the extracts as supported by the GC-MS analysis. Yet, only few studies have been performed on the MOSs extracts indicating the enhanced anti-cancerous potency of such extracts correlated to the occurrences of high oleic acid and fatty acid contents [50-52].

### 3.5. Anti-microbial efficacy of MOSs extracts

The Agar well diffusion assay was used to determine the anti-microbial effectiveness of the proposed MOSs extracts where the inhibited areas around the inoculated wells were measured. This zone of inhibition was caused by the diffusion of the active chemical constituents of the MOSs extract around the inoculated wells. The results revealed the considerable impact of the MOSs extracts seed on the bacterial strains (both gram-positive and gram-negative). However, the anti-bacterial action of the MOSs extracts was better against the gram-positive *S. aureus* bacteria. Fig 8 shows the MOSs extracts concentration dependent inhibition zones of the *S. aureus* bacteria (ranged from 18 to 24 mm). Conversely, the inhibition zones of the *E. coli* were ranged from 6 to 20 mm wherein the maximum and minimum inhibition zone diameter was obtained with the extracts concentration of 250 and 50 mg/ml, respectively (Fig 9). The observed discrepancy in the bactericidal action of the MOSs extracts for two different types of test bacteria may be due to the occurrence of their varying cell components [42].

**Fig 8.**
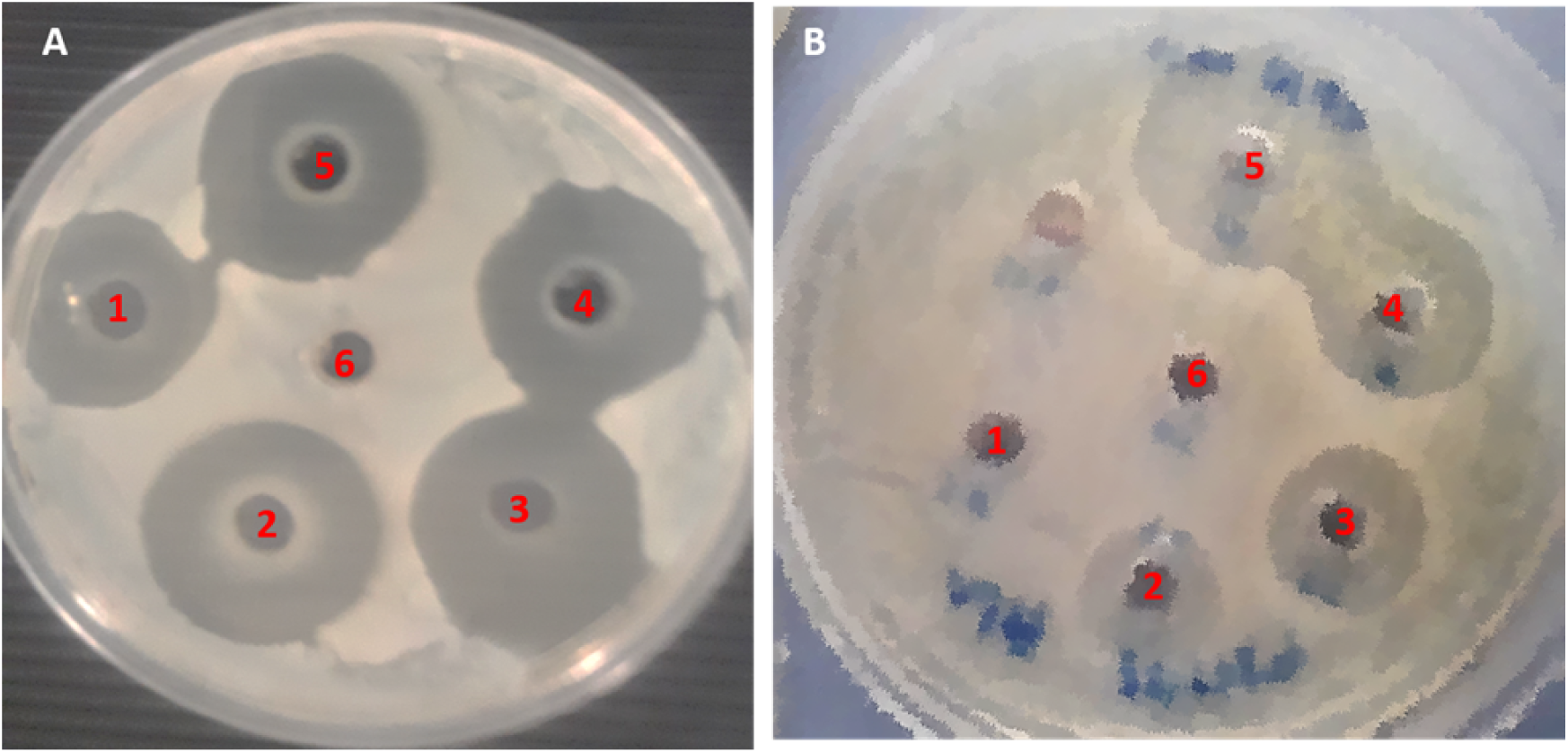
Agar well diffusion plates showing the zone of inhibition. (A) *S. aureus* (B) *E. coli.* **1**: 50 mg/ml, **2**: 100 g/ml, **3**: 150 mg/ml, **4**: 200 mg/ml, **5**: 250 mg/ml of seed extract of *Moringa oleifera*.

**Fig 9.**
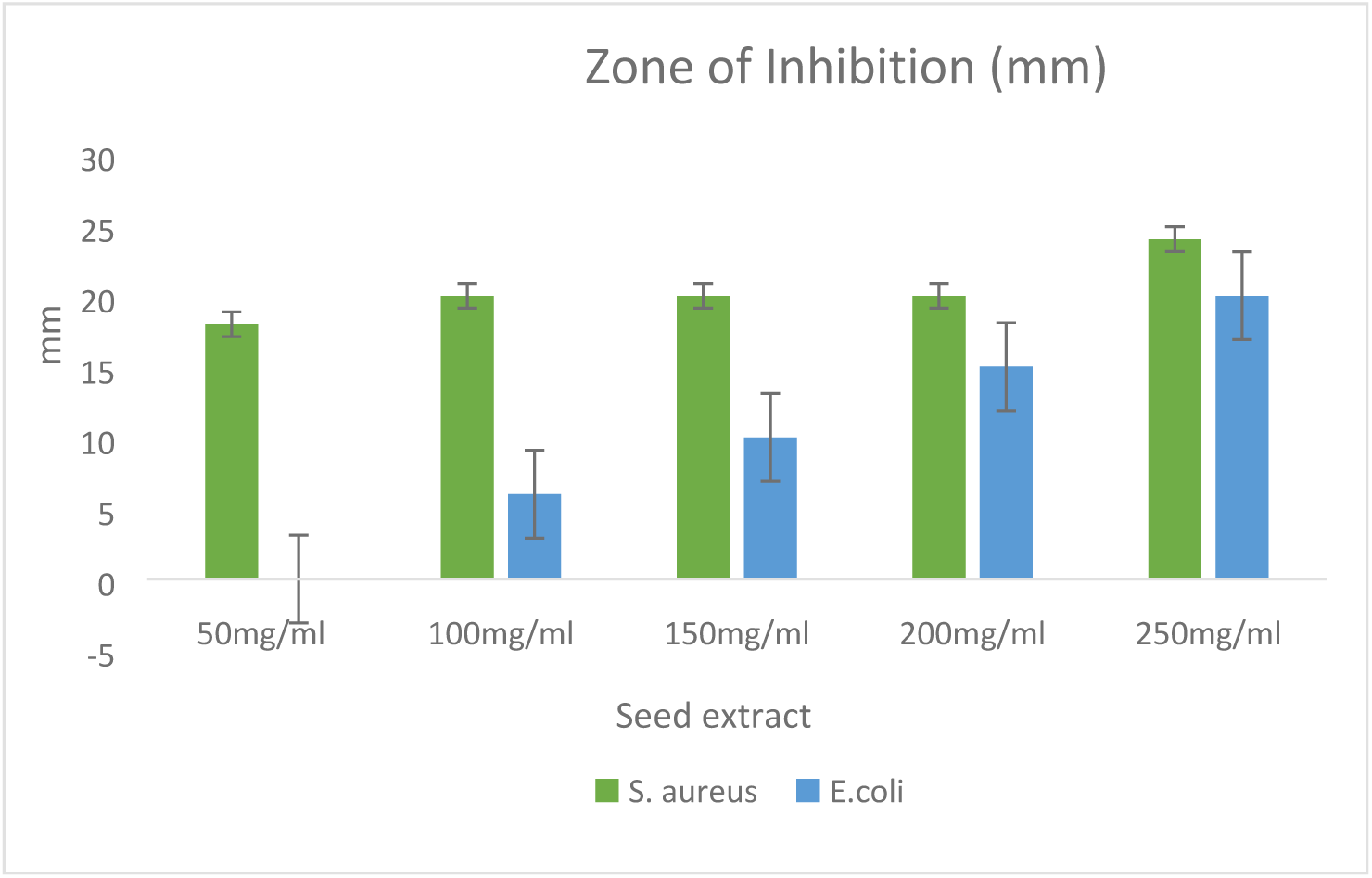
Zone of inhibition in millimeters (mm) against S. *aureus* and *E. coli* using the different concentrations of *Moringa oleifera* seed extract.

## 4. Conclusions

The LIBS and GC-MS technique was used to identify and quantify the elemental compositions of the MOSs. The anti-cancerous and anti-microbial effectiveness of the MOSs extracts were evaluated for the first time. The LIBS spectra revealed the presence of diverse elements in the MOSs useful for many health benefits. The GC-MS analysis reconfirmed the presence of several anti-cancerous and anti-microbial compounds in the MOSs extracts. The cell viability and DNA nuclear morphologies of the HEK-293 and HCT-116 cells treated with the MOSs extracts were measured using the MTT assay and DAPI staining, respectively. For the anti-cancer efficacy evaluation the HCT-116 were treated with MOSs extracts for 48 hrs before the test. The viability of the HEK-293 cells was examined after treating those using MOSs extracts. The cell viability percentage and IC_50_ values for both normal and cancerous cells were assessed. The MTT assay showed a significant impact of the MOSs extract for inhibiting the growth of the HCT-116 cells and insignificant inhibitory action of such extracts on the HEK-293 cells, indicating the excellent specificity of the extracts towards cancer cells. The MOSs extracts showed strong anti-microbial activity in terms of the growth inhibition when tested with *S. aureus* and *E. coli* bacteria using Agar well diffusion assay. It is established that MOSs extract can be a prospective anti-cancer and anti-microbial agent for the functional biomedicinal drug formulation.

## ^1^Abbreviations

MOSs: *Moringa oleifera* seed;
LIBS: Laser Induced Breakdown Spectroscopy;
GC-MS: Gas chromatography–mass spectrometry.

## Availability of data and material

The data generated during the current study are available from the corresponding author upon request.

## Author Declaration

We wish to confirm that there are no known conflicts of interest associated with this publication and there has been no significant financial support for this work that could have influenced its outcome.

## Authors’ Consent

All authors have approved the final version of the manuscript.

## Acknowledgements

Authors are thankful to the Deanship of Scientific Research (DSR) King Fahd University of Petroleum and Minerals and Institute for Research & Medical Consultations (IMRC), Imam Abdulrahman Bin Faisal University, Dammam, Saudi Arabia for providing the financial assistance [Project number 2019-015-IRMC] in carrying out the experiments. We also thank the help of Mr. Dionecio Jr. Bagon Dela Roca to carry out cell culture and bioassay. Authors are also grateful to Dr. Khaldoon M. Alsamman, Clinical Laboratory Science, College of Applied Medical Science, Imam Abdulrahman Bin Faisal University for providing HCT-116 and HEK-293 cell lines.

